# Cigarette smoking upregulates vascular expression of the novel atherosclerosis risk factor ADAMTS-7

**DOI:** 10.1101/2025.10.28.685237

**Authors:** Carla Abrahamian, M. Amin Sharifi, Tan An Dang, Aldo Moggio, Michael Winkler, Johannes Riechel, Hanna Winter, Baiba Vilne, Simone Kraut, Stefan Hadzic, Cheng-Yu Wu, Irina Pugach, Marija Gredic, Mirja Fassbender, Julia Hinterdobler, Julia Werner, Nikita Panyam, Moritz von Scheidt, Zouhair Aherrahrou, Yvonne Döring, Christian Graesser, Lars Maegdefessel, Norbert Weissmann, Heribert Schunkert, Hendrik B. Sager, Thorsten Kessler

**Affiliations:** Department of Cardiovascular Diseases, TUM University Hospital German Heart Center, Munich, Germany; German Centre for Cardiovascular Research (DZHK e.V.), partner site Munich Heart Alliance, Munich, Germany; Department of Internal Medicine III – Cardiology and Angiology, Saarland University Medical Center, Homburg/Saar, Germany; Vascular Biology and Experimental Vascular Medicine Unit, Department of Vascular and Endovascular Surgery, Klinikum rechts der Isar, School of Medicine and Health, Technical University Munich, Munich, Germany; Bioinformatics Group, Rīga Stradiņš University, Rīga, Latvia; Excellence Cluster Cardio-Pulmonary Institute (CPI), Institute for Lung Health (ILH) Universities of Giessen and Marburg Lung Center (UGMLC), Member of the German Center for Lung Research (DZL), Justus-Liebig-University, Giessen, Germany; Translational Animal Research Center (TARC), University Medicine at Johannes-Gutenberg University, Mainz, Germany; Institute for Cardiogenetics, University of Lübeck, Lübeck, Germany; German Centre for Cardiovascular Research (DZHK e.V.), partner site Hamburg/Kiel/Lübeck, Lübeck, Germany; Institute for Cardiovascular Prevention (IPEK), Ludwig Maximilian University, Munich, Germany; Department for BioMedical Research (DBMR), and Division of Angiology, Swiss Cardiovascular Center, Inselspital, Bern University Hospital, University of Bern, Bern, Switzerland

**Author notes:** Drs. Sager and Kessler contributed equally to this article as lead authors and supervised the work. Corresponding authors: Thorsten Kessler, MD Saarland University Medical Center Internal Medicine III – Cardiology and Angiology Saarland University Kirrberger Str. 100 · 66424 Homburg, Germany · Hendrik Sager, MD TUM University Hospital German Heart Center Department of Cardiovascular Diseases Technical University of Munich Lazarettstr. 36 · 80636 Munich, Germany ·. Drs. Abrahamian and Sharifi contributed equally to this article as first authors.

## Abstract

**Background:** Cigarette smoking is an established risk factor for coronary artery disease (CAD) and myocardial infarction. Genetic variants in the extracellular matrix protease ADAMTS-7 were also identified to increase CAD risk. Notably, *ADAMTS7* represents the only genomic locus that revealed a gene-environment interaction with smoking. The underlying mechanisms of this interaction remain unclear.

**Methods and Results:** In a murine model, cigarette smoke exposure (CSE) led to an upregulation of vascular *ADAMTS7* expression in wild type (WT) C57BL/6J mice. *ADAMTS7* upregulation was also found in carotid plaques from ever-smokers undergoing carotid endarterectomy in humans. Bulk RNA sequencing of lung tissues from WT mice exposed to CS revealed a downregulation of 20 and an upregulation of 173 transcripts. Among upregulated transcripts in smoking-exposed lungs, we found C-C motif chemokine ligand 17 (CCL17), which was likewise upregulated in plasma from smoking mice and humans. *In vitro*, recombinant CCL17 upregulated *ADAMTS7* expression in primary vascular smooth muscle cells (VSMC), which was inhibited secondary to silencing of CCL17’s *bona fide* receptor C-C Motif Chemokine Receptor 4 (CCR4). Conditioned media from CCL17-stimulated VSMC lacking ADAMTS-7 showed reduced release of inflammatory cytokines by endothelial cells (EC), reduced EC activation, and monocyte-to-EC adhesion. In proatherogenic *Apoe*^-/-^ mice exposed to CS, more numerous neutrophils, inflammatory monocytes, and macrophages were found in atherosclerotic plaques as compared to room air exposition. This effect was blunted in *Apoe*^-/-^*Adamts7*^-/-^ mice.

**Conclusions:** For the first time, our findings link CSE to vascular inflammation via CCL17-mediated upregulation of the CAD risk factor *ADAMTS7* and provide a mechanistic explanation for the gene-environment interaction between CS and *ADAMTS7* in CAD. Targeting ADAMTS-7 might be a promising therapeutic strategy irrespective of smoking status.

**Graphical Abstract:** 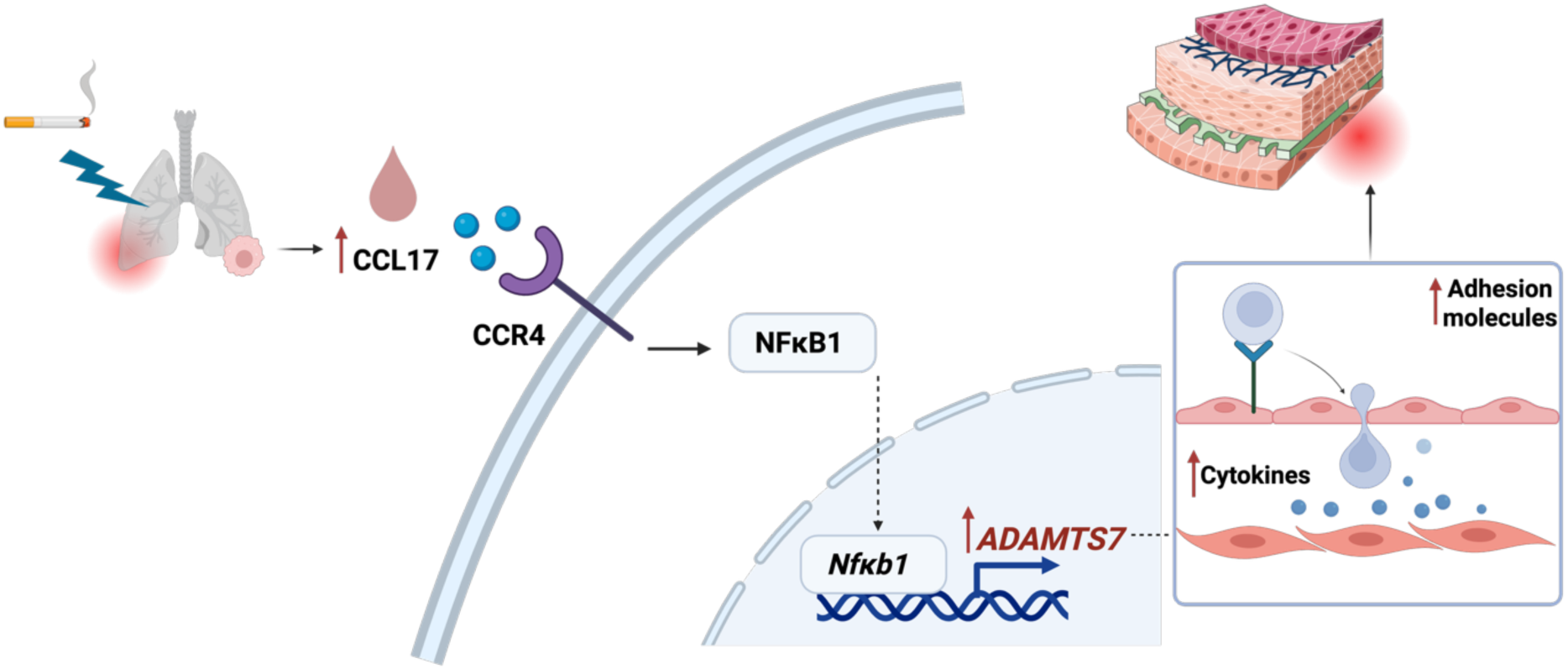

## Introduction

Ischemic heart disease remains the leading cause of morbidity and mortality worldwide^1^. Reducing its burden on individuals and healthcare systems is a major global health priority. Coronary artery disease (CAD) and myocardial infarction (MI), the primary manifestations of ischemic heart disease, are driven by a combination of risk factors. Recent analyses suggest that over 50% of CAD risk can be attributed to modifiable factors, including obesity, hypertension, diabetes, hypercholesterolemia, and cigarette smoking^2^.

Familial clustering of CAD and MI has long been recognized^3^, but it is only in the past two decades that genome-wide association studies (GWAS) have identified more than 300 loci associated with disease susceptibility^4^. One of the first loci identified harbors the *ADAMTS7* gene, encoding the extracellular matrix (ECM) protease ADAMTS-7^5–8^. Initially, its role in vascular biology was unclear. However, current evidence suggests that ADAMTS-7 contributes to vascular remodeling and atherosclerosis by degrading targets such as cartilage oligomeric matrix protein and thrombospondin-1, which impairs vascular healing and promotes neointima formation^9,10^. It also influences plaque formation and stability by degrading TIMP-1, an endogenous inhibitor of matrix metalloproteinases^11,12^. These properties make ADAMTS-7 a promising therapeutic target, as supported by recent studies exploring ADAMTS-7 vaccination in preclinical models^13^.

Beyond identifying genetic risk loci, GWAS have raised the prospect of uncovering gene-environment interactions to better explain interindividual variation in CAD risk. To date, however, only one robust interaction, i.e., between *ADAMTS7* and cigarette smoking, has been described. Genetic studies show that the protective allele at the *ADAMTS7* locus is associated with reduced risk of CAD risk in non-smokers, but this benefit is attenuated in smokers. *In vitro* exposure of vascular smooth muscle cells (VSMCs) – a key source of ADAMTS-7 in the vasculature – to cigarette smoke (CS) extract leads to increased *ADAMTS7* expression^14^. However, the complex composition of CS makes it difficult to identify the specific responsible mediators.

Despite strong genetic and epidemiological evidence linking both *ADAMTS7* and smoking to CAD, the mechanistic basis of their interaction remains unclear. Specifically, it is unknown (i) whether smoking directly upregulates *ADAMTS7* in the vasculature, (ii) which molecular mediators are involved, and (iii) what the functional consequences are. In this study, we investigated this gene-environment interaction using mouse models, *in vitro* experiments with primary human vascular cells, and analyses of human biospecimens.

## Methods

### Animal experiments

All animal experiments were conducted in accordance with German animal protection legislation and were approved by the local animal care committee (GI 20/10 Nr. G 57/2016). Mice of both sexes in a 1:1 ratio were used, housed under a 14/10 h light–dark cycle with *ad libitum* access to food and water. *Apoe^-/-^Adamts7^-/-^* (B6.*Adamts7^tm1e(KOMP)Wtsi^Apoe^tm1Unc^*) mice were generated by crossbreeding *Adamts7^-/-^* (B6.*Adamts7^tm1e(KOMP)Wtsi^*) with *Apoe^-/-^* (B6.129P2-*Apoe^tm1Un^*^c^) mice (purchased from Jackson Laboratories, Bar Harbor, ME, USA). Crossbreeding process was carried out for more than five generations as described previously^12^, and all experiments were performed using *Apoe^-/-^*and *Apoe^-/-^Adamts7^-/-^* littermates and C57BL6/J mice as control. Mice were maintained on either a chow or a Western diet (TD.88137, Teklad Harlan; Inotiv, Lafayette, IN, USA). The Western diet was started 3 days prior to cigarette smoke exposure (CSE) or in room air (RA) controls.

For CSE, mice were exposed to mainstream smoke from 3R4F cigarettes (Lexington, KY, USA) using a CS generator (Burkhart, Wedel, Germany). The exposure protocol was run for 6 h per day, 5 days per week, for 14 weeks at a concentration of 200 mg particulate matter per m^3^, following previously established protocols^15–17^. Age-matched control mice were kept under identical conditions but without CSE (room air, RA). Mice were randomly allocated to treatment and control groups, and all data from animal experiments were analysed in a blinded manner.

### ADAMTS7 expression in human carotid artery endarterectomy samples

Carotid atherosclerotic lesions were obtained from the Munich Vascular Biobank^18^, including samples from both smoking (active and ex-smokers) and non-smoking individuals. The biobank was approved by the local ethics committee (2799/10, Ethics Committee of the Faculty for Medicine at the Technical University of Munich, Munich, Germany) and is in accordance with the Declaration of Helsinki. The Illumina NovaSeq platform was used for library generation of extracted RNA of tissue biopsies taken during carotid endarterectomy using the miRNeasy Mini Kit (Qiagen, Hilden, Germany). RNA was extracted from total RNA using poly-T oligo-attached magnetic beads. Then, first-strand cDNA was generated with the TruSeq stranded total RNA kit and the TruSeq stranded mRNA kit (Illumina, San Diego, USA). Second strand cDNA was subsequently created with dUTP. End repair, adaptor ligation, size selection, A-tailing, amplification and purification were performed before the library was completed. Quality control was done using Qubit (Thermo Fisher, Waltham, USA), and real-time PCR as well as size distribution detection with the Bioanalyzer was performed (Agilent, Santa Clara, USA). The company Fios Genomics (Fios Genomics, Edinburgh, UK) evaluated the raw FASTQ data, which also included quality control and alignment using STAR aligner. Detailed methods were described previously^19,20^.

### CCL17 measurements in human plasma samples

Olink Explore HT plasma proteomics data from 525 coronary heart disease patients were matched to demographic records. Active smokers and ex-smokers were combined into one group. Propensity score matching was applied to balance never-smoker and smoker groups by age and sex using the MatchIt R package^21^. CCL17 plasma levels were compared between 217 smokers and 217 never-smokers.

### Transcriptome analysis of lung tissue

Total RNA was extracted from lung tissue of RA- or CS-exposed mice using the RNeasy Micro Kit according to the manufacturer’s protocol. RNA integrity and concentration were assessed by IMGM Laboratories GmbH (Martinsried, Germany) using the Agilent 2100 Bioanalyzer with the RNA 6000 Nano kit. Samples with an RNA Integrity Number ≥ 7 and DV200 ≥ 70% were considered suitable for sequencing. Quality control of sequencing data was performed by using CLC Genomics Workbench 12.0 on a total of 362 million reads across 10 datasets. Metrics such as GC content, base quality scores (PHRED), sequence duplication rates, and overrepresented k-mers were evaluated to confirm data quality.

Transcriptomic data from smoking and non-smoking mice were analyzed for differential gene expression using two independent statistical pipelines LIMMA (Linear Models for Microarray and RNA-Seq Data) and DESeq2. Genes were considered significantly differentially expressed if they met the thresholds of |log_2_ fold change| ≥ 1 and false discovery rate (FDR) < 0.05.

### Primary cell lines

Primary human Coronary Artery Smooth Muscle Cells (CASMCs) (C-14052; PromoCell, Heidelberg, Germany) were cultured in Smooth Muscle Cell Growth Medium 2 supplemented with 5% (v/v) fetal calf serum (FCS; C-22062, PromoCell) and 1% penicillin-streptomycin (10,000 U/mL) (15140122; Thermo Fisher, Waltham, MA, USA). CASMCs were washed with 1x phosphate-buffered saline (PBS) (14190-169; Thermo Fisher) and detached from the cell culture plates using 0.25% trypsin-EDTA (25200072; Thermo Fisher).

Human Umbilical Vein Endothelial Cells (HUVECs) (C-12203; PromoCell) were cultured in endothelial cell growth media supplemented with 2% (v/v) FCS (C-22110; PromoCell). HUVECs were washed with HEPES-buffered saline solution (C-40020; PromoCell) and detached using trypsin-EDTA (0.04%/0.03%; C-41020; PromoCell), followed by neutralization with trypsin neutralizing solution (C-41120; PromoCell).

The human monocyte cell line THP-1 (TIB-202; ATCC; Manassas, VA, USA) was cultured in RPMI 1640 Medium (A1049101; Thermo Fisher) supplemented with 10% fetal bovine serum (FBS; S0615; Sigma-Aldrich, St. Louis, MO, USA) and 1% penicillin-streptomycin (15140122; Thermo Fisher). All cells were maintained in a humidified incubator at 37 °C with 5% CO_2_.

### RNA interference and recombinant protein experiments

Human CASMCs were transfected with small interfering RNAs (siRNAs) using Lipofectamine RNAiMAX transfection reagent (13778150; Thermo Fisher) at approximately 75% confluence. Lyophilized siRNA (5 nmol) was reconstituted in DNase-/RNase-free water to a 10 μM working solution. Transfection reagents and siRNA were prepared following the manufacturer’s instructions. After transfection, cells were incubated in a humidified environment at 37 °C with 5% CO_2_ for up to 48 h before harvesting.

The following Silencer Select siRNAs (all from Thermo Fisher) were used: Silencer™ Select Negative Control (4390843), human *CCR4* (s3207), human *ADAMTS7* (s22052), and human *NFKB1* (s9504). Knockdown efficiency was determined by quantitative real-time polymerase chain reaction (qPCR), as detailed below.

For recombinant protein stimulation, CASMCs were treated with human CCL17 (ab283903; Abcam, Cambridge, UK) at 100 nM and/or 500 nM (as indicated in the respective figure legends) for 24 h, similarly for all: human IL-6 (206-IL-010; R&D Systems, Minneapolis, MN, USA), human CXCL1 (275-GR; R&D Systems), and human CCL2 (279-MC; R&D Systems).

### Preparation of mouse tissues

For flow cytometry analysis of aortic leukocytes, aortas were excised from the aortic arch near the heart down to the iliac bifurcation in the lower abdomen, carefully removing perivascular fat and surrounding tissues. Isolated aortas were then finely minced and subjected to enzymatic digestion in a buffer containing collagenase I (450 U/ml), collagenase XI (125 U/ml), DNase I (60 U/ml), and hyaluronidase (60 U/ml) (all from Sigma-Aldrich). Digestion was carried out at 37 °C with continuous agitation at 750 rpm for 1 h using a thermoshaker. Lung samples were processed using the same digestion protocol. Following digestion, cell suspensions were passed through a 40 µm cell strainer to remove debris, centrifuged, and resuspended to obtain single-cell suspensions for further analysis.

### Blood sampling

For flow cytometry of peripheral blood leukocytes, blood samples were incubated with 1x red blood cell (RBC) lysis buffer (BioLegend, San Diego, CA, USA) for 5 min to remove red blood cells. The reaction was stopped, and cells were resuspended in FACS buffer (PBS) containing 0.5% bovine serum albumin (Sigma-Aldrich) to maintain cell stability prior to analysis. For blood chemistry and other analyses, blood samples were collected in EDTA-coated microvettes, kept on ice, and centrifuged at 1,000 g for 15 min at 4 °C. The resulting plasma was used for biochemical measurements or enzyme-linked immunosorbent assays.

### Flow cytometry

All stainings were performed at 4°C in 300 µl of FACS buffer. Gating strategies were applied based on forward scatter (FSC-A) and side scatter (SSC-A), adjusted for tissue type to identify viable cells (FSC-A vs. SSC-A) and single cells (FSC-A vs. FSC-W and SSC-A vs. SSC-W). Total leukocytes and specific subsets were distinguished using a hematopoietic lineage cocktail (B220, CD90.2, CD49b, NK1.1, Ter-119, and Ly6G) along with CD45.2, CD11b, F4/80, and Ly6C. Details on fluorochromes are provided in **Suppl. Table S1**.

Neutrophils were defined as lineage^high^CD45.2^high^CD11b^high^. Monocytes were defined as lineage CD45.2^high^CD11b^high^lineage^low^Ly6C^high^ F4/80^intermediate^. Macrophages were defined as lineage CD45.2^high^CD11b^high^lineage^low^Ly6C^high^F4/80^high^. In the lung, interstitial macrophages were identified as MHCll^high^CD11b^high^, and alveolar macrophages MHCll^high^CD11b^intermediate^.

Compensation was performed using OneComp eBeads (01-1111-42; Thermo Fisher) conjugated with the respective antibodies. Flow cytometry data were acquired on an LSRFortessa (BD Biosciences, Franklin Lakes, NJ, USA) and analyzed using FlowJo software version 10 (BD Biosciences).

### Immunoblotting

Cells were washed with Dulbecco’s phosphate-buffered saline (PBS; Biochrom, Berlin, Germany) and collected. Cell lysis was performed using radioimmunoprecipitation assay (RIPA) buffer (Sigma-Aldrich). To enhance membrane disruption, samples were subjected to sonication (3 x 30 s), with 30 s intervals on ice between each cycle. Lysates were clarified by centrifugation, and supernatants were collected for protein analysis. Prior to electrophoresis, samples were mixed 1:1 ratio with 2x Laemmli buffer (Sigma-Aldrich) and boiled at 95°C for 5 min.

Protein samples (10-20 μg per lane) were separated by SDS-PAGE on 4–20% pre-cast gradient gels (Bio-Rad, Hercules, USA) and transferred onto methanol-activated polyvinylidene difluoride (PVDF) membranes (Bio-Rad) at 300 V for 20 min. Membranes were blocked with either 5% dry milk (A0830.1000; VWR, Radnor, PA, USA) or bovine serum albumin (BSA; A1391,0500; Applichem, Darmstadt, Germany) in PBS for 1 h at room temperature before overnight incubation at 4 °C with primary antibodies prepared in 2.5% dry milk or 5% BSA.

Following incubation, membranes were washed with PBS-T (PBS containing 0.1% v/v Tween 20; Sigma-Aldrich) and incubated for 1 h at room temperature with HRP-conjugated secondary antibodies (anti-mouse or anti-rabbit IgG; Cell Signaling Technology, Danvers, MA, USA) at 1:10,000 dilution. After additional washes with PBS-T, protein bands were detected using an enhanced chemiluminescence detection system (GE Healthcare Life Sciences, Chicago, IL, USA) and imaged with the ImageQuant 800 system (Amersham Biosciences, Amersham, UK). Band intensity quantification was performed using ImageJ (RRID: SCR_003070). GAPDH or Vinculin were used as loading controls. Antibody dilutions and buffer compositions are listed in **Suppl. Table S1**.

### Quantitative real-time polymerase chain reaction

Total RNA was extracted from cell lines using the RNeasy Plus Mini Kit (74134; Qiagen, Hilden, Germany) according to the manufacturer’s protocol. Complimentary DNA (cDNA) was synthesized from total RNA using the RevertAid First Strand cDNA Synthesis Kit (K1622; Thermo Fisher). Relative mRNA expression levels were assessed via quantitative real-time PCR (qPCR) using the PerfeCTa SYBR Green FastMix (low ROX; Quantabio, Beverly, MA, USA) according to the manufacturer’s instructions. qPCR was performed using the Applied Biosystems ViiA 7 Real-Time PCR System with the ViiA 7 software v.1.2.2 (Thermo Fisher). Relative target gene expression was calculated by normalization to the reference gene *GAPDH*. Primer sequences are provided in **Suppl. Table S2**.

### Enzyme-linked immunosorbent assays (ELISA)

Plasma samples from mice or human cell culture supernatants were collected as previously described. Plasma samples were diluted 1:5 and supernatants 1:2 using the corresponding assay dilution buffers. Cytokine levels in plasma were measured using CCL17 (ab213474; Abcam) and IL-6 (ab100712; Abcam) ELISA kits. In cell culture supernatants, GM-CSF (DGM00), IL-1β (DLB50), CCL7 (DCC700), VEGF (DVE00), HGF (DHG00B), ICAM1 (DCD540), and CCL20 (DM3A00; all from R&D Systems) were used to quantify cytokine concentrations. All assays were performed according to the manufacturers’ instructions, and absorbance was measured at 450 nm with background correction at 540 nm using the Infinite M200 PRO microplate reader (Tecan, Maennedorf, Switzerland). Final concentrations were calculated from a standard curve and four-parameter logistic (4PL) regression.

### Cytokine profiling

As described above, human CASMCs were transfected with either non-silencing control siRNA (siRNA_scramble_) or ADAMTS-7 siRNA (siRNA*_ADAMTS7_*) for 48 h. On the following day, the medium was replaced with serum-free medium, either supplemented with or without CCL17, and the cells were incubated for an additional 24 h under low-volume conditions to concentrate the treatment.

Following stimulation with CCL17, the Biotechne Proteome Profiler Human Cytokine Array Kit (ARY022B; R&D Systems) was used to analyze cytokine expression in CASMCs supernatants. Membranes were blocked and then incubated with CASMCs supernatant overnight. The next day, membranes were washed three times using 1x Wash Buffer, followed by incubation with detection antibodies, and 1x Streptavidin-HRP, separated by wash cycles. Following chemiluminescent development, cytokine signals were visualized with Amersham ImageQuant 800 system Scanner (Cytiva, using the integrated IQ800 software).

Each membrane included six reference spots at three corners, used for alignment with a transparency overlay template and serving as internal positive controls. Cytokine spot signals were visualized using ImageQuant TL Array version 8.1, and pixel intensities of each cytokine spot were quantified. Duplicate spot signals for each analyte were averaged, background-subtracted using the signal from a clear region or negative control, and normalized to the average intensity of the six reference spots. A cut-off value of 0.05 for the lowest normalized signal intensity in the siRNA_scramble_+CCL17 group was applied to exclude analytes with signals below this threshold from further analysis. Cytokines with a relative difference ≥ ±50% were considered for validation using ELISA.

### Leukocyte adhesion assay

Human CASMCs were seeded to 70% confluence and transfected with either non-silencing negative control siRNA or ADAMTS-7 siRNA for 48 h, as described above. On the following day, the medium was replaced with serum-free medium with or without CCL17 for an additional 24 h. Meanwhile, HUVECs were seeded into gelatin-coated 48-well plates and cultured for 48 h until a confluent monolayer formed. HUVECs were then incubated with conditioned medium from CASMCs cultures for 24 h. THP-1 cells were harvested at 1·10^6^ cells/mL in serum-free medium and labeled with 1× LeukoTracker (Cell Biolabs, San Diego, CA, USA) for 1 h at 37 °C. Cells were washed twice with serum-free medium. HUVEC monolayers were washed with serum-free medium, and 200 μL of the labeled THP-1 cell suspension was added to each well. After 90 min of incubation, non-adherent cells were removed by gentle washing with wash buffer.

For fluorescence-based quantification, adherent cells were lysed with lysis buffer for 5 min on a shaker, lysates were transferred to 96-well plates, and fluorescence was measured at 480 nm excitation/520 nm emission using an Infinite M200 PRO microplate reader (Tecan). For confocal imaging, cells were fixed in 4% paraformaldehyde (15714-1L; Electron Microscopy Sciences, Hatfield, PA, USA) for 20 min at room temperature, washed twice with PBS, and blocked with 5% BSA in PBS for 45 min. Cells were stained with anti-CD31 overnight at 4 °C, followed by incubation with Alexa Fluor 594–conjugated secondary antibody and DAPI for 1.5 h at room temperature. After two washes with PBS, adherent leukocytes were visualized using the confocal laser scanning microscope (STELLARIS 5; Leica Microsystems, Wetzlar, Germany). For z-stack imaging, sequential optical sections were acquired across the full depth of the endothelial monolayer using a 63×/1.4 NA oil immersion objective. Optical sections were collected at 0.3–0.5 μm intervals, covering the entire monolayer from the basal to the apical surface. THP-1 cells were visualized via LeukoTracker (488 nm; Cell Biolabs), endothelial cells with Alexa Fluor 594 (594 nm; Thermo Fisher), and nuclei with DAPI (405 nm; Thermo Fisher). Maximum-intensity projections and 3D reconstructions were generated from the z-stacks using LAS X software (Leica Microsystems).

### Statistical analysis

Data are presented as mean ± standard error of the mean (s.e.m.), median and interquartile range (IQR), or violin plots, as appropriate. Outliers were identified and removed using the ROUT method. Bulk transcriptome analyses were performed as described above. Otherwise, distribution of data was analyzed using D’Agostino & Pearson omnibus test or Shapiro-Wilk test, as appropriate. Normally distributed data were analyzed using unpaired t-test for two comparisons or one-way ANOVA for more than two comparisons with appropriate post-hoc tests for multiple comparisons (depicted in the figure legends). Non-normally distributed data were analyzed using Mann-Whitney test for two comparisons or Kruskal-Wallis test for more than two comparisons with appropriate post-hoc tests for multiple comparisons (depicted in the figure legends). All tests were two-sided. Results of distribution analyses, the applied statistical tests, and all experimental details are provided in **Supplemental Table S3**. P-values/adjusted p-values <0.05 were considered statistically significant. GraphPad Prism for macOS, version 10.4.2 (GraphPad, Boston, MA, USA), was used for most analyses unless otherwise stated.

## Results

### Smoking upregulates vascular ADAMTS7 expression in humans and mice

We first sought to investigate the influence of smoking on Adamts7 expression in murine vascular tissues. To that end, WT mice were exposed to cigarette smoke (CS) as compared to RA for 98 days (**Fig. 1A**). When we analyzed *Adamts7* mRNA levels in aorta using qPCR, we found that CSE led to an upregulation of *Adamts7* as compared to RA exposure (1.41±0.16, n=15, vs. 1.04±0.08 [2^-τιCt^], n=15, p=4.75·10^-2^; **Fig. 1B**). On protein level, the difference was more pronounced with higher Adamts-7 protein levels in mice that were exposed to CS (157.0±17.3, n=5, vs. 38.0±14.2 [a.u.], n=5, p=7.1·10^-4^; **Fig. 1C**). To test whether smoking similarly affects vascular *ADAMTS7* expression in humans, we probed plaque specimen from individuals undergoing carotid endarterectomy which were included in the Munich Vascular Biobank. In smokers, plaque *ADAMTS7* mRNA levels were higher as compared to non-smokers (median 5.72 [interquartile range (IQR) 5.34-5.96], n=71, vs. 5.43 [IQR 4.92-5.8] [a.u.], n=104, p=2.21·10^-3^; **Fig. 1D**). Taken together, these data indicate that exposure to CS upregulates vascular ADAMTS-7 production.

**Figure 1:**
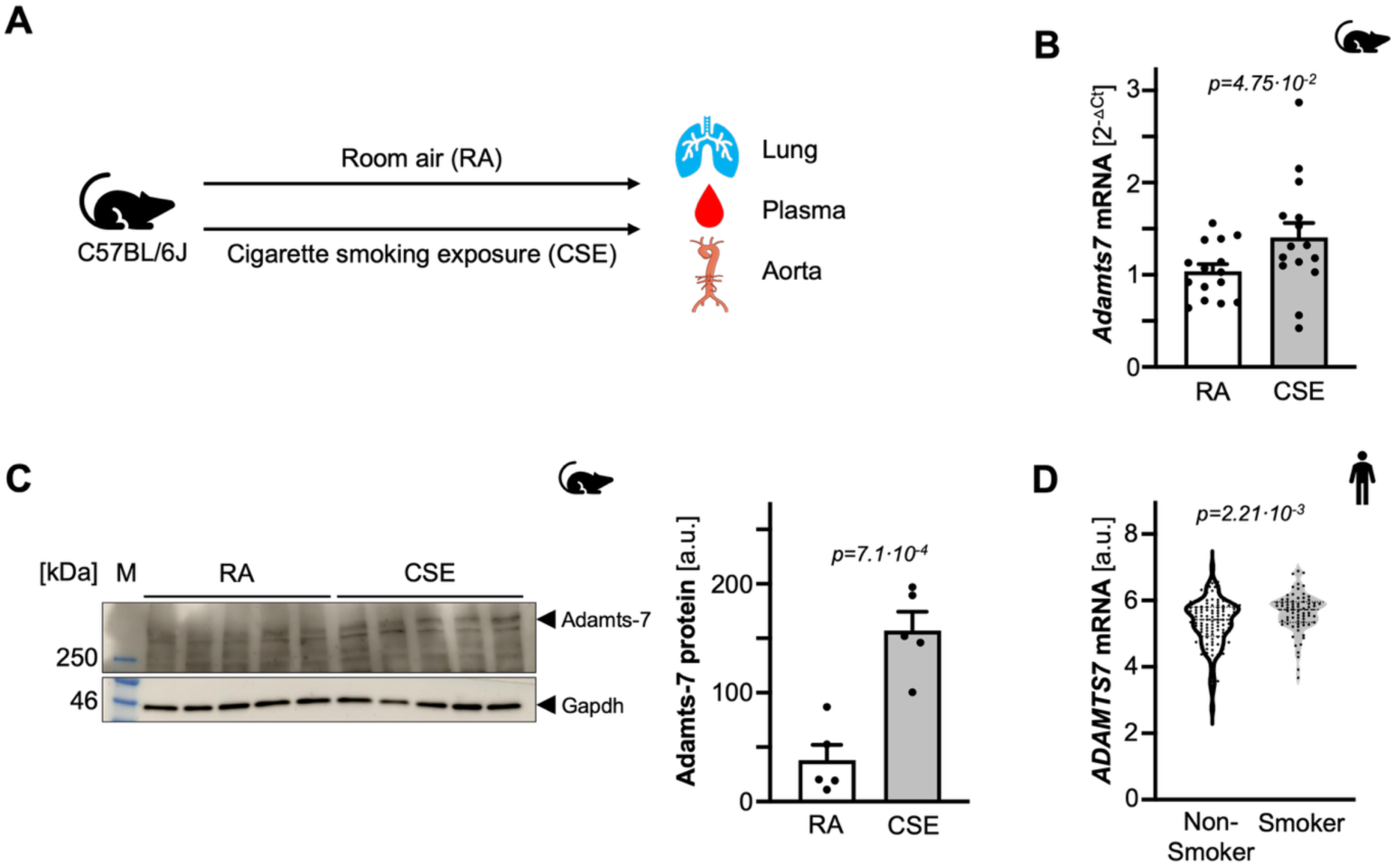
Smoking upregulates ADAMTS-7 in vascular tissues. **A**. Schematic overview. **B**. *Adamts7* expression in murine aorta samples secondary to room air or cigarette smoke exposure (n=15 each). **C**. Left: representative immunoblot of aorta samples from individual mice exposed to room air or cigarette smoke. Right: quantification of protein levels (n=5 each). **D**. *ADAMTS7* expression in human carotid plaque specimen of non-smokers and smokers (n= 104 vs. 71). Each symbol indicates one individual/animal. Data are mean and s.e.m. (**B**, **C**) or median and quartiles (**D**). Unpaired t-test (**B**, **C**) or Mann-Whitney test (**D**). Abbreviations: *a.u.*, arbitrary units; *CSE*, cigarette smoke exposure; *RA*, room air.

### Pulmonary inflammation following cigarette smoking exposure

The lung represents the first organ which is exposed to the toxic effects of cigarette smoking. Therefore, we asked whether changes in *ADAMTS7* expression in vascular tissue might be a consequence of a prior CS-related lung injury. To assess pulmonary immune cell composition, we performed flow cytometry gating of lung single-cell suspensions (**Fig. 2A**). First, we exposed WT mice to either room air or CS for 98 days. Flow cytometry analysis showed that CSE led to an increase of interstitial macrophages (33.37±3.51 vs. 16.02±2.14, n=13 vs. 15, p=3.97·10^-4^; **Fig. 2B**) whereas alveolar macrophages were reduced (24.04±2.05 vs. 60.58±1.33, n=13 vs. 15, p=5.99·10^-14^; **Fig. 2C**) in mice exposed to CS as compared to RA, consistent with chronic lung inflammation. We then performed transcriptome analysis of whole lung tissue to unravel dysregulated inflammatory pathways, visualized in a volcano plot (**Fig. 2D**). Out of 48,526 transcripts, we found 193 to be differentially expressed between the conditions. Of these, 173 were up- and 20 were downregulated (**Suppl. Table S4**). **Table 1** displays the 15 most strongly upregulated transcripts in mice exposed to CS. These included genes encoding chemoattractant proteins such as Saa3, as well as matrix metalloproteases and inflammatory cytokines. These data indicate that CSE leads to local inflammation in the lung and the production of inflammatory cytokines with putative systemic effects.

**Figure 2:**
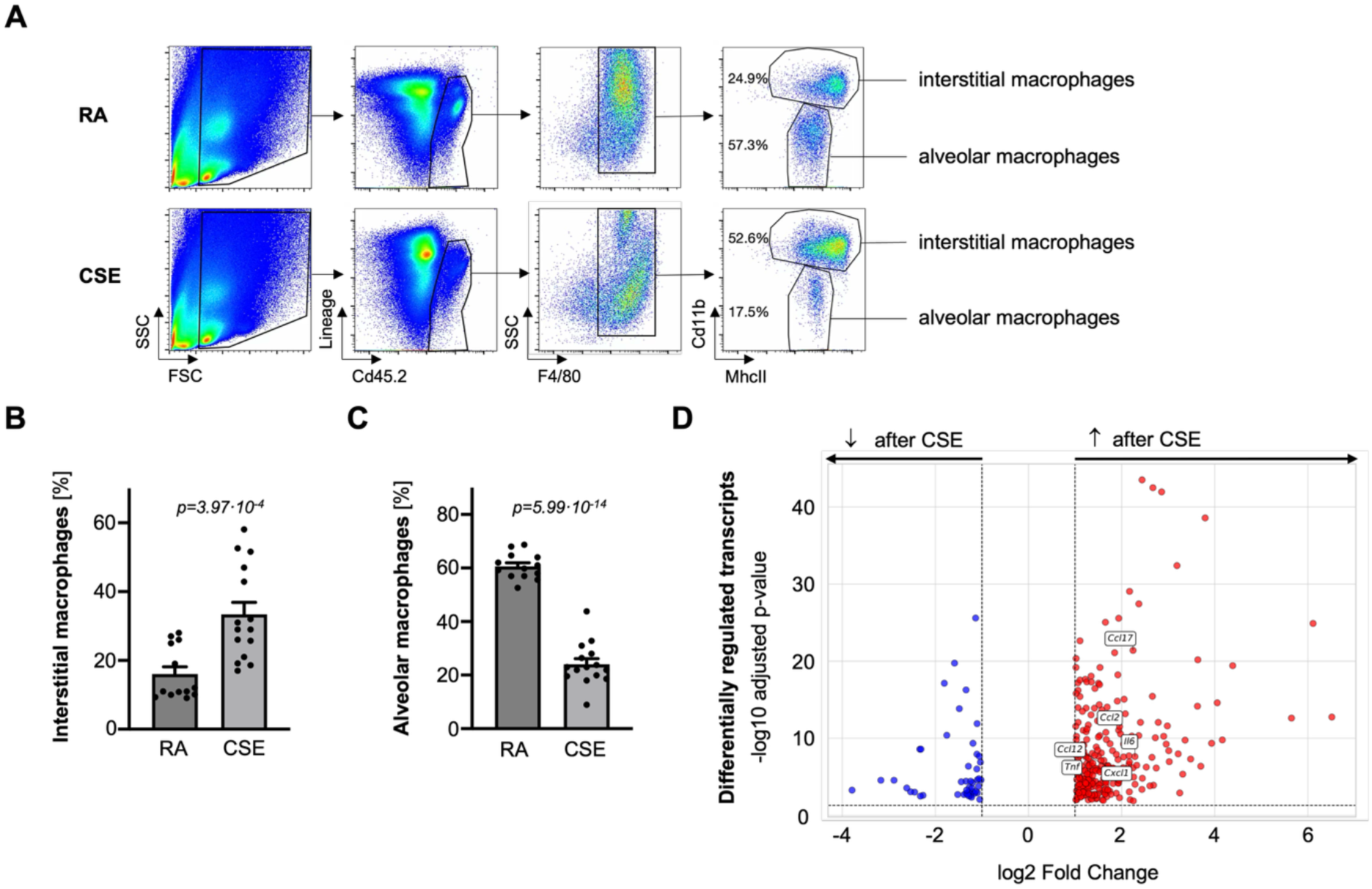
Pulmonary inflammation secondary to cigarette smoking. **A**. Gating strategy to quantify interstitial and alveolar macrophages in lung tissue of mice which were subjected to room air or cigarette smoke exposure. **B**, **C**. Proportions of interstitial (**B**) and alveolar macrophages (**C**) in mice exposed to room air or cigarette smoke (n= 13 vs. 15). **D**. RNA sequencing of lung tissue samples displaying up- (red) and downregulated (blue) transcripts secondary to cigarette smoking as compared to room air exposure. Each symbol indicates one animal (**B**, **C**) or transcript (**D**). Data are mean and s.e.m. (**B**, **C**). Unpaired t-test (**B**, **C**) or Benjamini–Hochberg procedure (**D**). Abbreviations: *CSE*, cigarette smoke exposure; *RA*, room air.

**Table 1:** Top 15 upregulated transcripts in wild type mice which were exposed to cigarette smoking for 98 days as compared to room air. P-values are adjusted for multiple testing accoding to the Benjamini-Hochberg procedure.

| Gene | Function/description | Mean log2 fold-change | p-value |
| --- | --- | --- | --- |
| *Rph3a* | Neurotransmitter release, strongly associated with smoking status | 6.51 | $1.14 \cdot 10^{-13}$ |
| *Saa3* | chemoattractant activity, secreted | 4.05 | $1.62 \cdot 10^{-15}$ |
| *Cyp1a1* | high expression in lung, correlated to smoking status, associated with lung cancer risk | 3.79 | $1.75 \cdot 10^{-39}$ |
| *Cyp2a4* | correlation to smoking in GWAS | 3.49 | $3.20 \cdot 10^{-8}$ |
| *Cd177* | neutrophil activation | 3.19 | $2.66 \cdot 10^{-33}$ |
| *Mmp12* | pulmonary emphysema | 2.96 | $3.7 \cdot 10^{-10}$ |
| *Arnt2* | hypoxia response | 2.67 | $2.14 \cdot 10^{-43}$ |
| *Cxcr1* | interleukin 8 receptor, neutrophil chemotaxis | 2.40 | $5.63 \cdot 10^{-13}$ |
| *Il6* | pro-inflammatory cytokine | 2.28 | $1.56 \cdot 10^{-8}$ |
| *Ccl17* | T cell chemotaxis | 2.24 | $2.61 \cdot 10^{-22}$ |
| *Cxcl1* | neutrophil attractant | 2.16 | $1.60 \cdot 10^{-7}$ |
| *Ctsk* | degradation of extracellular matrix, associated with monocyte count | 2.05 | $5.62 \cdot 10^{-16}$ |
| *Ccl2* | monocyte attractant | 1.96 | $2 \cdot 10^{-12}$ |
| *Ccl12* | monocyte active chemokine, important role during early allergic lung inflammation | 1.66 | $2.61 \cdot 10^{-7}$ |
| *Timp1* | inhibition of MMPs | 1.58 | $5.05 \cdot 10^{-8}$ |

### CCL17 upregulates ADAMTS7 in vascular smooth muscle cells

In mouse lung tissue, we found five cytokines among the top upregulated transcripts: interleukin 6 (*Il6*), C-X-C motif chemokine ligand 1 (*Cxcl1*), and chemokine (C-C) ligands 2 (*Ccl2*), 12 (*Ccl12*), and 17 (*Ccl17*). We decided to not further investigate *Ccl12* as it is not conserved in humans. Using a cytokine profiler, we next investigated plasma levels of Il6 and Ccl17. While Il6 plasma levels were comparable between mice exposed to cigarette smoking and room air (**Suppl. Fig. S1)**, Ccl17 plasma levels were elevated by 70% secondary to smoking exposure (1.7±0.1 vs. 1.0±0.09 [fold-change], n=4, p=1.70·10^-3^; **Fig. 3A**). Importantly, also in human smokers, we found higher levels of plasma CCL17 compared to never-smokers (median [interquartile range, IQR]: 0.06 [-0.38–0.77] vs. -0.13 [-0.69–0.39] [normalized protein expression], n=217 each, p=8.72·10^-4^; **Fig. 3B**). Cxcl1 and Ccl2 were not present on the cytokine profiler. We thus next investigated the influence of these two cytokines and CCL17 on endogenous *ADAMTS7* expression in primary human coronary artery smooth muscle cells (CASMCs). CXCL1, IL6 and CCL2 did not alter *ADAMTS7* expression (**Suppl. Fig. S2)**. In contrast, incubation of CASMCs with CCL17 upregulated endogenous *ADAMTS7* expression by almost 36% (135.7±15.1 vs. 100±5.5 [%], n=11 each, p=3.79·10^-2^; **Fig. 3C**).

**Figure 3:**
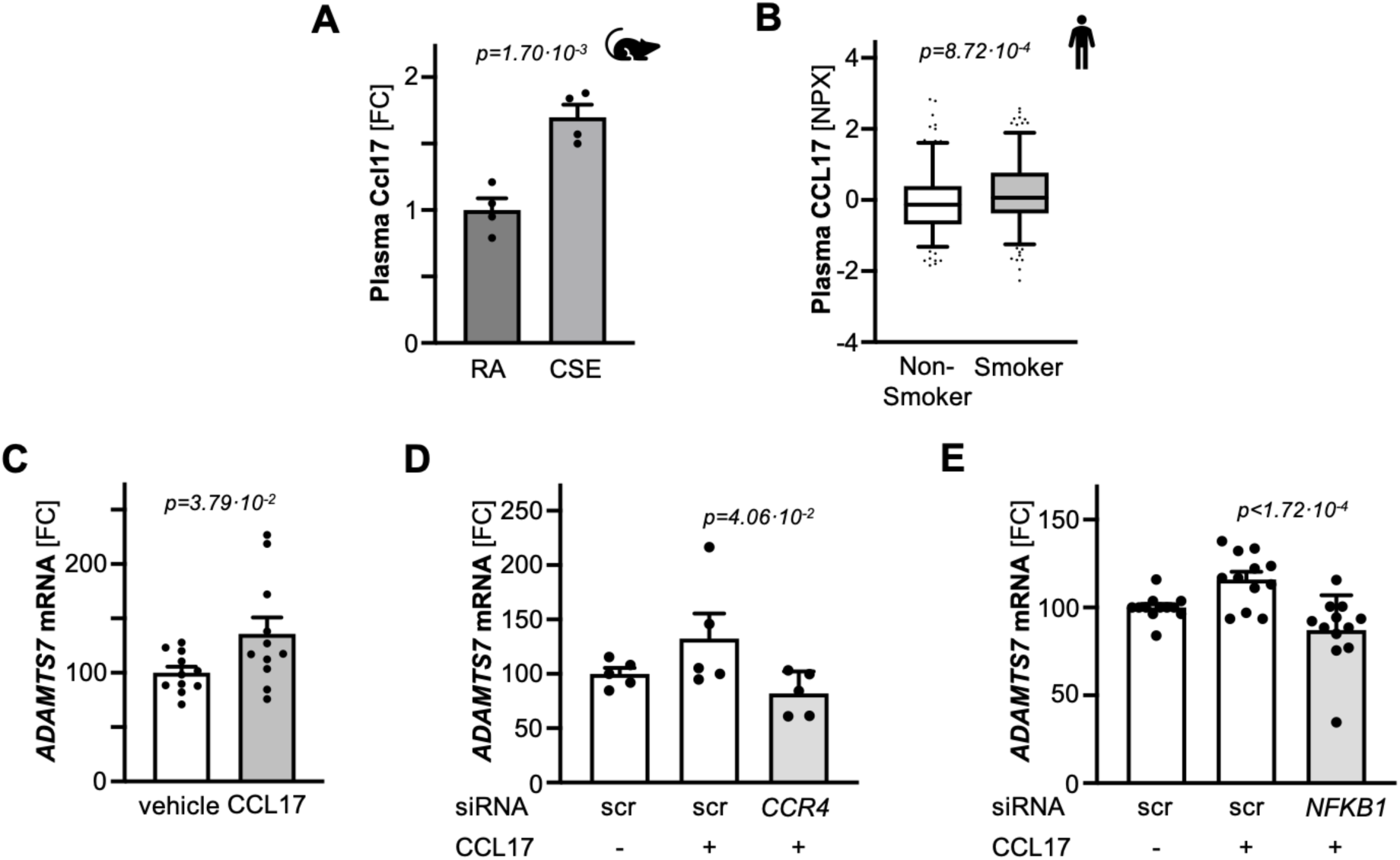
CCL17 influences ADAMTS-7 levels in vascular smooth muscle cells. **A**. Plasma Ccl17 levels in mice which were exposed to room air or cigarette smoke (n=4 each). **B**. CCL17 plasma levels in human non-smokers and smokers (n= 217 each). **C**. Expression of *ADAMTS7* in vascular smooth muscle cells which were incubated with 500 ng CCL17 (n=11 each). **D**, **E**. Expression of *ADAMTS7* in vascular smooth muscle cells secondary to incubation with vehicle, CCL17 and/or knockdown of *CCR4* (n= 5 each) (**D**) or *NFKB1* (n= 12 each) (**E**) compared to control (scr) using RNA interference. Each symbol indicates one animal/individual (**A**, **B**) or independent experiment (**C**-**E**). Data are mean and s.e.m. (**A**, **C**-**E**) or median and quartiles (**B**). Unpaired t-test (**A**, **C**), Mann-Whitney test (**B**) and ordinary one-way ANOVA with Šídák’s test for multiple comparisons (**D**, **E**). Abbreviations: *CCL17*, C-C motif chemokine 17; *CSE*, cigarette smoke exposure; *FC*, fold-change; *NPX*, normalized protein expression; *scr*, scramble.

CCL17 was initially described as a thymus-regulated and T-cell chemokine cytokine with a pivotal function in (auto-) inflammatory diseases and in cancer^22^. In a publicly available single cell RNA sequencing dataset from mice with chronic CSE^23^, we found that monocytes and interstitial but also alveolar macrophages are the main source of Ccl17 (**Suppl. Fig. S3**). Its *bona fide* receptor, i.e., C-C chemokine receptor-4 (CCR4), is expressed in CASMCs on mRNA and protein level (**Suppl. Fig. S4)**. To investigate whether the upregulation of *ADAMTS7* is mediated via CCR4 signaling, we used RNA interference (RNAi) to downregulate *CCR4*. Using a small interfering RNA (siRNA) targeting *CCR4* (siRNA*_CCR4_*), endogenous *CCR4* expression was substantially reduced (**Suppl. Fig. S5A**). As a consequence, *ADAMTS7* expression secondary to CCL17 exposure was reduced as compared to scramble control (81.92±9.05 vs. 132.5±22.88 [%], n=5 each, p=4.06·10^-2^; **Fig. 3D**). Downstream CCR4 signaling involves the nuclear factor ’kappa-light-chain-enhancer’ of activated B-cells (NF-κB) pathway^24^. This pathway is also involved in tumor necrosis factor α (TNFα) signaling, which was previously described to modulate *ADAMTS7* expression^9^ and was also upregulated in lung tissue after smoking (**Table 1**). To test whether CCL17 impacts *ADAMTS7* expression through NF-κB signaling, we next silenced *NFKB1* expression in CASMCs using RNAi (**Suppl. Fig. S5B)**. Downregulation of *NFKB1* resulted in reduced expression of *ADAMTS7* secondary to CCL17 exposure as compared to scramble control (87.18±5.71 vs. 115.9±4.39 [%], n=12 each, p=1.72·10-4; **Fig. 3E**) In summary, these data indicate that CCL17 regulates endogenous *ADAMTS7* expression in vascular smooth muscle cells via the CCR4-NF-κB axis.

### ADAMTS-7 influences the release of inflammatory cytokines by vascular smooth muscle cells

The mechanisms by which ADAMTS-7 impacts atherosclerosis remain incompletely understood. To investigate whether the absence of ADAMTS-7 renders the secretome of vascular smooth muscle cells less inflammatory, we stimulated CASMCs with CCL17 following RNAi-mediated silencing of *ADAMTS7*. Endogenous *ADAMTS7* levels was reduced by 65% (**Suppl. Fig. S6)**. Explorative cytokine profiler analyses revealed a differential pattern of cytokine expression (**Fig. 4A**). Only highly expressed cytokines were considered for further analysis, applying a cut-off value of 0.05 for the lowest normalized intensity in the control group. Cytokines demonstrating a ≥ ±50% change in abundance were followed up using ELISA. Among these, Granulocyte-Macrophage Colony-Stimulating Factor (GM-CSF; 0.58±0.05 vs. 1.00±0.10, n=6, p=4.56·10^-3^), Hepatocyte Growth Factor (HGF; 0.31±0.03 vs. 1.00±0.08, n=6, p<1.15·10^-5^), Intracellular Adhesion Molecule 1 (ICAM1; 0.80±0.07 vs. 1.00±0.06, n=6, p=0.08), Interleukin 1 beta (IL-1 β ; 0.77±0.10 vs. 1.00±0.051, n=6, p=0.09), Monocyte Chemotactic Protein 3 (MCP-3; 0.41±0.04 vs. 1.00±0.09, n=6, p=1.53·10^-4^), Interleukin 1 beta (IL-1β; 0.77±0.10 vs. 1.00±0.051, n=6, p=0.09), C-C motif chemokine ligand 20 (CCL20; 0.87±0.18 vs. 1.00±0.09, n=6, p=0.64), and Vascular Endothelial Growth Factor (VEGF; 0.69±0.03 vs. 1.00±0.05, n=6, p=3.12·10^-4^), were detected at lower levels in supernatant from CCL17-treated CASMCs lacking ADAMTS-7 (**Fig. 4B**). These findings indicate that the lack of ADAMTS-7 attenuates the release of key cytokines implicated in leukocyte recruitment and vascular remodeling, thus modulating the pro-inflammatory secretome of vascular smooth muscle cells.

**Figure 4:**
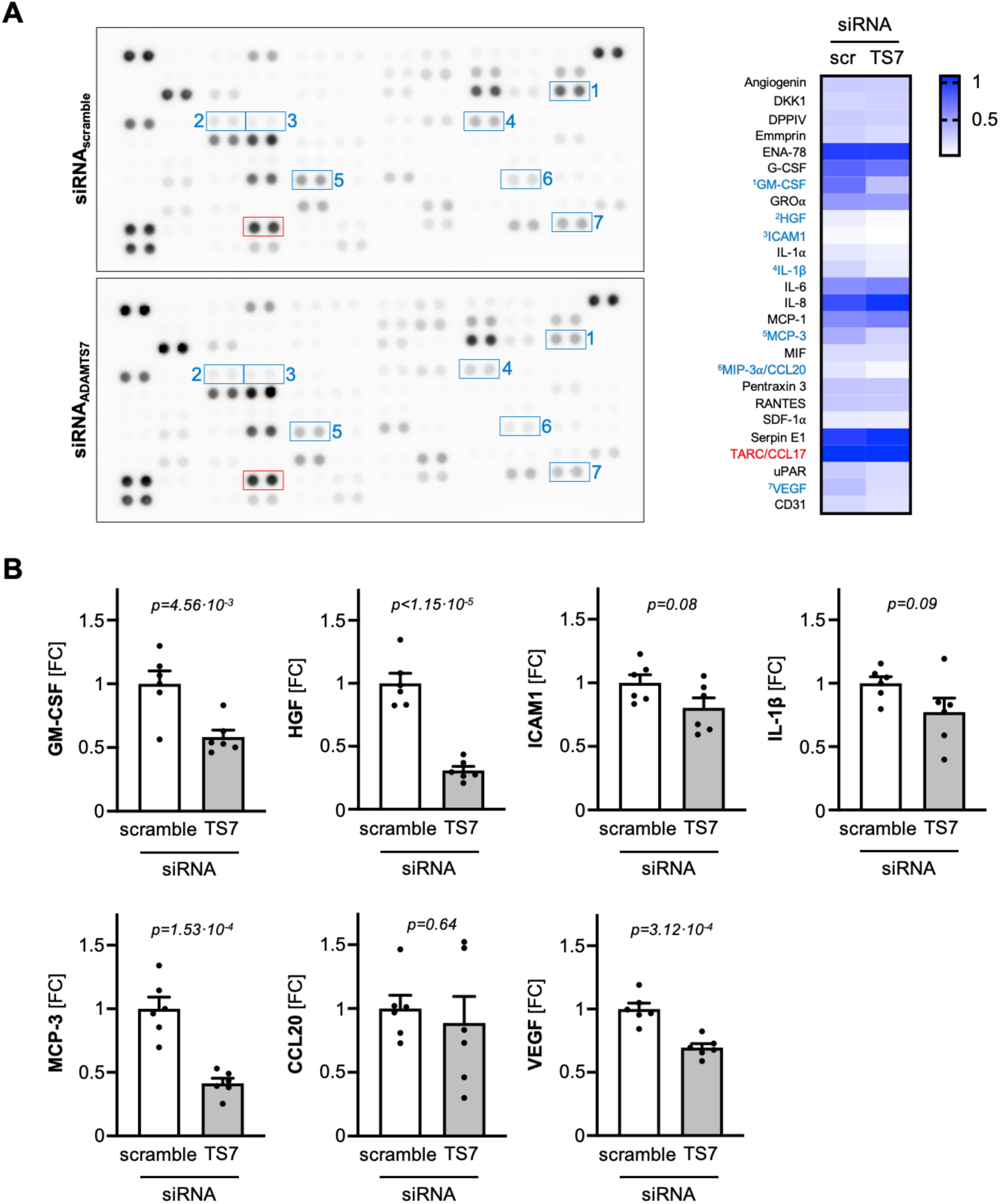
Impact of ADAMTS-7 deficiency on vascular smooth muscle cell supernatants secondary to incubation with CCL17. **A**. Representative cytokine profiler blots (left) and heatmap visualizing cytokine profiler results. Cytokines displaying a more than 50% difference between the conditions are highlighted in blue. CCL17 was similar between the groups (red). **B**. Validation of findings from A using enzyme-linked immunosorbent assays (n=6 each). Each symbol indicates one independent experiment. Data are mean and s.e.m. Unpaired t-test. Abbreviations: *CCL20*, C-C motif chemokine 20; *FC*, fold-change; *GM-CSF*, Granulocyte-Macrophage Colony-Stimulating Factor; HGF, Hepatocyte Growth Factor; *ICAM1*, intercellular adhesion molecule 1; *IL-1β*, Interleukin 1-β; *MCP*-*3*, Monocyte Chemotactic Protein 3; *scr*, scramble; *TS7*, ADAMTS-7; *VEGF*, Vascular Endothelial Growth Factor.

### VSMC-derived ADAMTS-7 influences endothelial cell activation

The recruitment of inflammatory leukocytes to the vascular wall is a hallmark of plaque initiation and progression^25^. Endothelial cells (EC) represent the barrier between circulating leukocytes and the vessel wall. Given that CCL17 altered the vascular smooth muscle cell secretome in an ADAMTS-7-depedent manner, we next investigated whether these changes may influence EC phenotypes. To that end, we generated conditioned media by incubating CASMCs with CCL17 in the absence and presence of ADAMTS-7 using RNAi. The supernatants were then transferred to ECs (**Fig. 5A**). We first analyzed the expression of a panel of adhesion molecules (**Fig. 5B**). As compared to conditioned media from control cells, conditioned media lacking ADAMTS-7 reduced the expression of transcripts encoding vascular cell adhesion molecule 1 (*VCAM1*; 31.61±11.14 vs. 100.00±9.98 [%], n=7 each, p=6.40·10^-4^), intercellular adhesion molecule 1 (*ICAM1*; 47.53±14.45 vs. 100.00±8.88, n=7 each, p=9.3·10^-3^), intercellular adhesion molecule 2 (*ICAM2*; 57.76±15.80 vs. 100.00±6.54, n=7, p=4.95·10^-2^), vascular endothelial cadherin (*CDH5*; 44.34±13.03 vs. 100.00±7.96, n=7, p=3.36·10^-3^), and E-selectin (*SELE*; 20.30±10.74 vs. 100.00±9.05, n=7, p=1.17·10^-3^). Expression levels of transcripts encoding platelet and endothelial cell adhesion molecule 1 (PECAM1) were not changed (**Suppl. Fig. S7)**.

**Figure 5:**
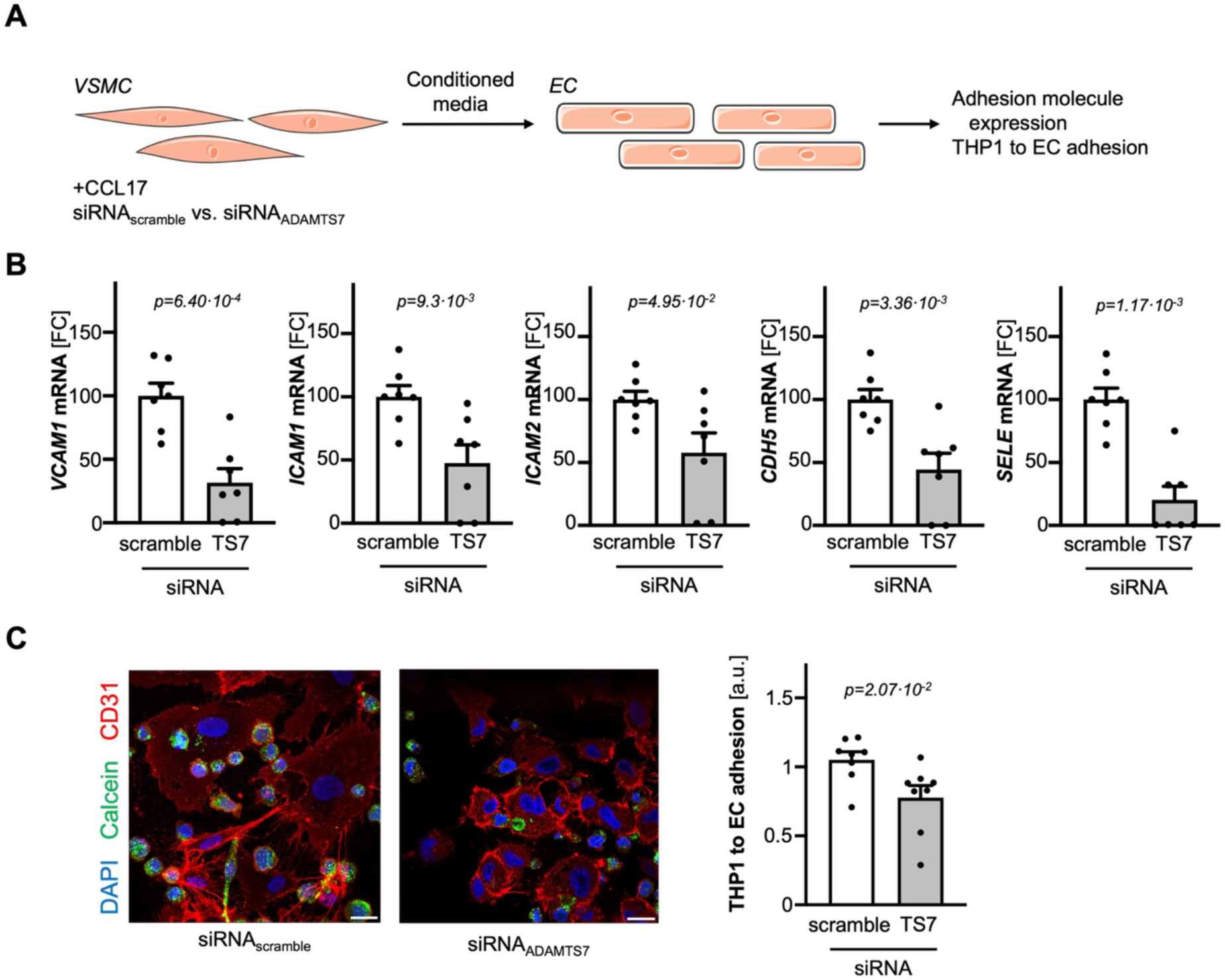
Reduced activation of endothelial cells by conditioned media of vascular smooth muscle cells lacking ADAMTS-7. **A**. Schematic overview of the experimental setup. **B**. Significantly downregulated adhesion molecules in the absence of ADAMTS-7 (n=7 each). **C**. Reduced adhesion of THP-1 cells to endothelial cells in the absence of ADAMTS-7 (n=8 each). Each symbol indicates one independent experiment. Data are mean and s.e.m. Unpaired t-test. Abbreviations: *CDH5*, VE-cadherin; *FC*, fold-change; *EC*, endothelial cell(s); *ICAM1*, intercellular adhesion molecule 1; *ICAM2*, intercellular adhesion molecule 2; *SELE*, E-selectin; *TS7*, ADAMTS-7; *VCAM1*, vascular cell adhesion molecule 1; *VSMC*, vascular smooth muscle cell(s).

We next aimed at investigating whether these molecular changes translate into functional differences in leukocyte to EC adhesion. Therefore, we incubated THP-1 cells (i.e., a human monocyte cell line) with EC which were pretreated with conditioned media as described above. Absence of ADAMTS-7 in CASMCs resulted in reduced adhesion of THP-1 cells to EC (0.78±0.09 vs. 1.05±0.06 [a.u.], n=8, p=2.07·10^-2^; **Fig. 5C**).

Taken together, our findings indicate that ADAMTS-7-dependent secretome of VSMCs modulates EC phenotypes, regulating the expression of adhesion molecules and promoting leukocyte adhesion, potentially contributing to inflammatory cell recruitment during atherogenesis.

### Lack of ADAMTS-7 reduces smoking-induced vascular inflammation in vivo

ADAMTS-7 has been implicated in atherosclerotic plaque formation. Our findings suggested that ADAMTS-7 may also influence vascular inflammation. To test this, we first investigated whether CS exposure affects vascular leukocyte numbers inside the vessel wall under proatherogenic conditions. We hence exposed *Apoe*^-/-^ mice, which were fed Western diet for 14 weeks, to either CS or RA and subsequently analyzed atherosclerotic aortas using flow cytometry. To determine whether ADAMTS-7 influences vascular inflammation, we included *Apoe*^-/-^*Adamts7*^-/-^ mice as a third group, also fed a Western diet and exposed to CS (**Fig. 6A**).

**Figure 6:**
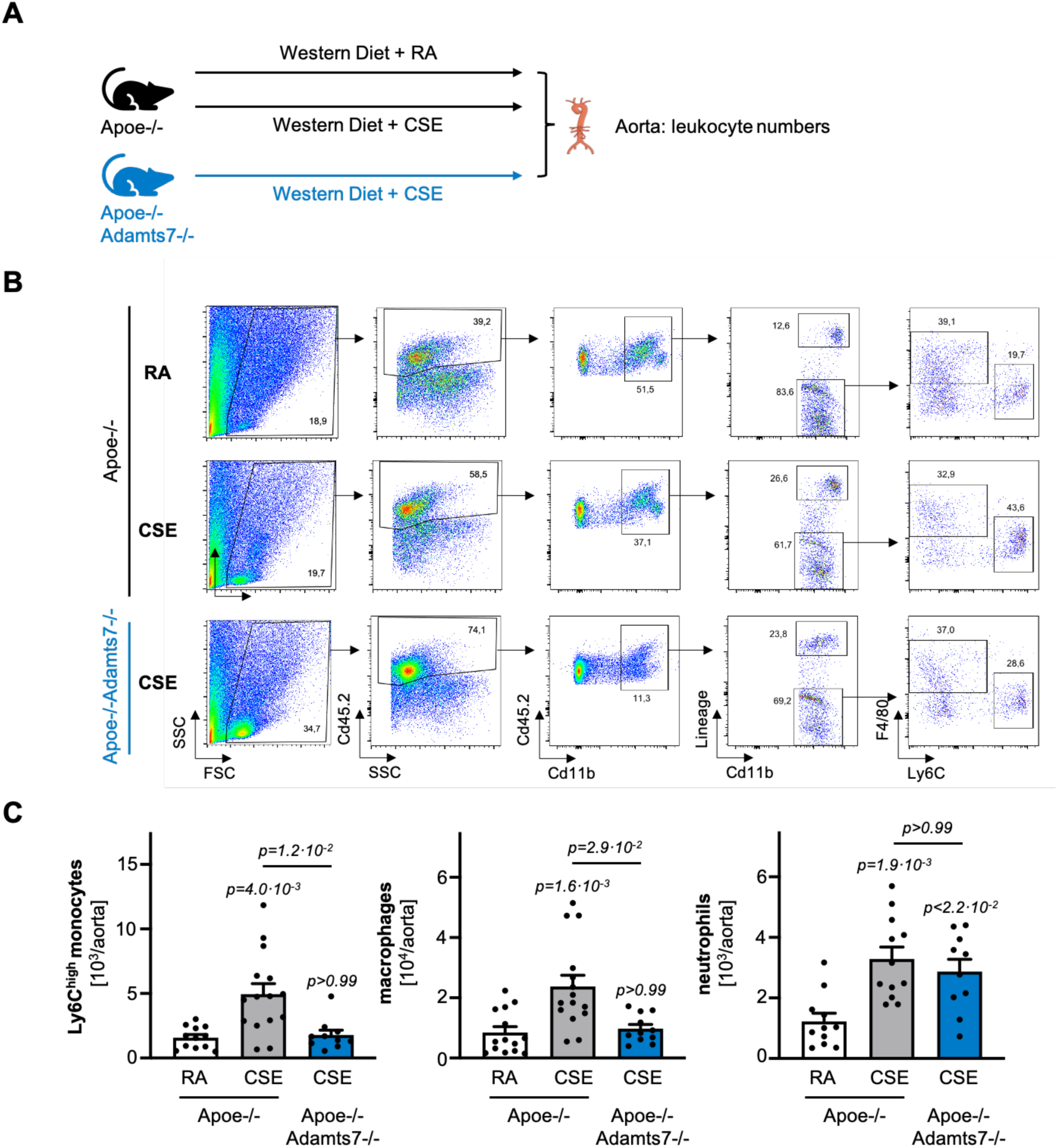
Influence of smoking and Adamts7 on vascular inflammation under proatherogenic conditions. **A**. Schematic overview of the experimental setup. **B**. Gating strategy to quantify Ly6Chigh monocytes, macrophages, and neutrophils in atherosclerotic aortas. **C**. Numbers of Ly6Chigh monocytes (left), macrophages (middle), and neutrophils (right) in samples from *Apoe*^-/-^ mice exposed to room air (n=11-15 as indicated above) or cigarette smoke (CSE; n=12-15 as indicated above), and *Apoe*^-/-^*Adamts7*^-/-^ mice exposed to cigarette smoke (CSE; n=10-11 as indicated above). Each symbol indicates one animal. Data are mean and s.e.m. One-way ANOVA with Šídák’s test for multiple comparisons (Ly6Chigh monocytes, neutrophils) or Kruskal-Wallis test, with Dunn’s test for multiple comparisons (macrophages). Abbreviations: *CSE*, cigarette smoke exposure; *RA*, room air.

In *Apoe*^-/-^ mice, CSE compared to RA resulted in significantly higher numbers of plaque neutrophils (3.28±0.39.2·10^3^/aorta, n=12, vs. 1.22±0.26·10^3^/aorta, n=11, p=1.9·10^-3^), Ly6C^high^ monocytes (4.95±0.80·10^3^/aorta, n=15, vs. 1.57±0.24·10^3^/aorta, n=12, p=4.0·10^-3^), and macrophages (2.37±0.37·10^4^/aorta, n=14 vs. 0.84±0.19·10^4^/aorta, n=15, p=1.6·10^-3^), indicating that smoking increased vascular inflammation under these conditions (**Fig. 6C**). In contrast, *Apoe*^-/-^*Adamts7*^-/-^ mice exposed to CS displayed lower numbers of Ly6C^high^ monocytes (1.78±0.37·10^3^/aorta, n=10, p>0.99 vs. Apoe^-/-^ RA, p=1.2·10^-2^ vs. Apoe^-/-^ CSE) and macrophages (9.72±0.14·10^3^/aorta, n=11, p>0.99 vs. *Apoe*^-/-^ RA, p=2.96·10^-2^ vs. *Apoe*^-/-^ CSE) as compared to *Apoe*^-/-^ mice exposed to CS and comparable numbers to *Apoe*^-/-^ mice exposed to RA. Numbers of plaque neutrophils were comparable between *Apoe*^-/-^ mice exposed to CS independent of ADAMTS-7 (2,870±407.0, n=10, p<2.2·10^-2^ vs. Apoe^-/-^ RA, p>0.99 vs. Apoe^-/-^ CSE; **Fig. 6B**, **C**). These data indicate that the absence of ADAMTS-7 *in vivo* partially attenuates vascular inflammation induced by CS.

## Discussion

Cigarette smoking is a major risk factor for cardiovascular disease and all-cause death^2,26^, promoting endothelial dysfunction, oxidative stress, and leukocyte recruitment^27,28^. Despite successful smoking cessation initiatives, the number of active smokers globally has remained stable at approximately 1.2 billion smokers^29^, underscoring its ongoing relevance. Here, using human and murine biospecimens, we show that cigarette smoking upregulates vascular *ADAMTS7* expression and identify CCL17 as a novel regulator of *ADAMTS7*. We furthermore report that absence of ADAMTS-7 beneficially influences vascular inflammation, particularly secondary to CS exposure.

Genome-wide association studies have reproducibly identified *ADAMTS7* as a CAD risk locus^5–7^. Gene-smoking analyses revealed that the protective *ADAMTS7* allele regarding CAD risk reduces CAD risk by ∼12%, but that this effect is blunted in smokers^14^ . *In vitro* experiments using CS extracts suggested higher expression of *ADAMTS7* in vascular smooth muscle cells, indicating a direct effect of one of the many unknown CS constituents on *ADAMTS7*’s regulatory circuit. Whether smoking indeed affects *ADAMTS7* expression remained speculative.

In a series of experiments, we found that CS exposure increased *ADAMTS7* mRNA levels in mouse aorta and human atherosclerotic carotid plaques. Importantly, in mice Adamts-7 protein levels were also elevated in aortic tissue secondary to smoke exposure. Together, these findings provide the first *in vivo* experimental evidence that cigarette smoking induces increased expression of a CAD risk gene in the vasculature.

Although an upregulation of *ADAMTS7* expression by CS constituents has been shown *in vitro*^14^ and remains biologically plausible, we wondered whether lung inflammation might remotely contribute to its regulation *in vivo*. In lung tissue from mice chronically exposed to CS, we observed a shift towards higher numbers of interstitial macrophages and increased expression of inflammatory transcripts. Among these, CCL17 emerged as a particularly promising candidate, as we found higher levels in plasma samples of both CS-exposed mice and human individuals with a history of smoking. Consistently, CCL17 was previously reported as one of the most strongly upregulated transcripts in CS exposure models^30^.

Our data suggest a spillover of CCL17 from the lung to the circulation, raising the question whether CCL17 influences *ADAMTS7* expression. The main receptors for CCL17 are CCR4 and CCR8, classically expressed on CD4+ T-cells^31,32^. Here, we show that CCR4 in contrast to CCR8 is also expressed in CASMCs, suggesting that CCL17 may exert direct effects via its canonical signaling pathway in resident vascular cells. Indeed, incubation of CASMCs with CCL17 led to a marked increase in *ADAMTS7* expression which was blunted by silencing of CCR4. The downstream signaling cascade involves NF-κB-dependent transcriptional mechanisms. Here, activation by CCL17 converges with other inflammatory pathways which upregulate *ADAMTS7* expression, e.g., TNFα^33^, but CCL17 has not been described to regulate ADAMTS-7 before. In our CS exposure experiment, we also identified TNFα transcripts to be upregulated in the lung. It therefore seems likely that a combination of inflammatory cytokines, including CCL17, contributes to the regulation of vascular *ADAMTS7* expression while CCL17 alone is able to substantially increase *ADAMTS7* expression.

While previous experimental studies have shown that lack of ADAMTS-7 is beneficial for neointima formation^10^ and atherosclerotic plaque phenotypes^11,12^, and that a vaccination against ADAMTS-7 is able to exert comparable effects^13^, details on molecular and cellular mechanisms involving ADAMTS-7 remain scarce. As an ECM protease, the known mechanisms involve heterocellular signaling with EC^10^ and leukocytes^12^ . Here, we demonstrate that lack of ADAMTS-7 leads to a less inflammatory profile of the supernatant of CASMCs *in vitro*. This, in turn, affects EC activation, resulting in reduced adhesion molecule expression and leukocyte-to-EC adhesion *in vitro*. While the exact mechanisms by which ADAMTS-7 modulates the inflammatory profile of VSMCs remains to be elucidated, reduced release of, e.g., MCP-3 provides a plausible explanation for the observed effects on leukocyte adhesion^34^ . Notably, we also observed increased numbers of leukocytes in the vascular wall of CS-as compared to RA-exposed *Apoe*^-/-^ mice. This effect was abrogated in *Apoe*^-/-^ mice exposed to CS but lacking Adamts-7, specifically for monocytes and macrophages. These findings confirm that the absence of ADAMTS-7 reduces vascular inflammation *in vivo*. It, however, remains unclear why neutrophil numbers were not affected by the absence of Adamts-7.

Our data confirm a role for ADAMTS-7 in atherosclerosis and vascular inflammation. Among the genetic loci identified by genome-wide association studies, ADAMTS-7 emerges as one of the few reproducible and mechanistically validated targets that may help reduce residual atherosclerotic risk despite optimized medical therapy. The interaction of ADAMTS-7 with smoking, in addition, seems important from at least three perspectives: (i) increased expression of *ADAMTS7* secondary to smoking is a concrete mechanistic link underscoring the detrimental vascular effects of smoking; (ii) the involvement of remote injury in the lung affecting vascular *ADAMTS7* expression highlights the systemic impact of smoking; (iii) an inhibition of ADAMTS-*7* may provide therapeutic benefit in reducing vascular inflammation in non-smokers and smokers.

Our study has several limitations. First, experiments in mice do not fully resemble the human organism. To mitigate this, we validated key findings – as increased ADAMTS7 expression after CS exposure and changes in CCL17 plasma levels – in human biospecimens. Second, we investigated only a single duration of CS exposure. The results might differ with alternative CS exposure protocols. In addition, the effects of smoking cessation on ADAMTS7 expression remain unknown. The observation that human biospecimens from both active and ever-smokers showed increased vascular *ADAMTS7* expression suggests a potential persistence of this effect, but this remains speculative. Third, the mechanistic evidence for a role of ADAMTS-7 in regulating EC activity is derived from cell culture experiments, which by nature cannot fully mimic *in vivo* conditions. Nevertheless, given prior evidence supporting a role for ADAMTS-7 in modulating EC phenotypes, we believe that this represents a biologically relevant mechanism. Finally, due to the combined burden of CS exposure and Western diet in our animal model, we were not able to disentangle leukocyte recruitment as the underlying mechanism of different leukocyte numbers in the vascular walls of *Apoe*^-/-^ and *Apoe*^-/-^ *Adamts7*^-/-^ mice.

Taken together, we identified ADAMTS-7 as a molecular link between cigarette smoking and vascular inflammation. By uncovering that the CCL17-CCR4-NF-κB axis drives *ADAMTS7* expression, we provide mechanistic insight into how cigarette smoking modulates vascular risk. These findings furthermore highlight ADAMTS-7 as a promising therapeutic target to reduce residual cardiovascular risk, particularly in individuals with a history of smoking.

## Data Availability

The authors declare that all supporting data are available within the article and its Online Supplementary Files.

## Acknowledgments

The authors thank Christopher Wolf for technical assistance. **Figure 5A** contains modified image material available at Servier Medical Art under a Creative Commons Attribution 3.0 Unported License. The **Graphical Abstract** was created with BioRender.com. Generative AI (ChatGPT-4o, OpenAI) was used for language refinement and proofreading.

